# A new lineage of segmented RNA viruses infecting animals

**DOI:** 10.1101/741645

**Authors:** Darren J. Obbard, Mang Shi, Katherine E. Roberts, Ben Longdon, Alice B. Dennis

## Abstract

Metagenomic sequencing has revolutionised our knowledge of virus diversity, with new virus sequences being reported faster than ever before. However, virus discovery from metagenomic sequencing usually depends on detectable homology: without a sufficiently close relative, so-called ‘dark’ virus sequences remain unrecognisable. An alternative approach is to use virus-identification methods that do not depend on detecting homology, such as virus recognition by host antiviral immunity. For example, virus-derived small RNAs have previously been used to propose ‘dark’ virus sequences associated with the Drosophilidae (Diptera). Here we combine published *Drosophila* data with a comprehensive search of transcriptomic sequences and selected meta-transcriptomic datasets to identify a completely new lineage of segmented positive-sense single-stranded RNA viruses that we provisionally refer to as the *Quenyaviruses*. Each of the five segments contains a single open reading frame, with most encoding proteins showing no detectable similarity to characterised viruses, and one sharing a small number of residues with the RNA-dependent RNA polymerases of single- and double-stranded RNA viruses. Using these sequences, we identify close relatives in approximately 20 arthropods, including insects, crustaceans, spiders and a myriapod. Using a more conserved sequence from the putative polymerase, we further identify relatives in meta-transcriptomic datasets from gut, gill, and lung tissues of vertebrates, reflecting infections of vertebrates or of their associated parasites. Our data illustrate the utility of small RNAs to detect viruses with limited sequence conservation, and provide robust evidence for a new deeply divergent and phylogenetically distinct RNA virus lineage.

## 1 Introduction

Pioneered by studies of oceanic phage (Breitbart et al. 2002), since the mid-2000s metagenomic studies have identified thousands of new viruses (or virus-like sequences) associated with bacteria, plants, animals, fungi, and single-celled eukaryotes (reviewed in Greninger 2018, Obbard 2018, Shi et al. 2018, Zhang et al. 2018). At the same time, routine high-throughput sequencing has provided a rich resource for virus discovery among eukaryotic host genomes and transcriptomes (e.g. Bekal et al. 2011, Longdon et al. 2015, Webster et al. 2015, François et al. 2016, Mushegian et al. 2016, Gilbert et al. 2019). Indeed, a recent survey suggested that, as of 2018, around 10% of the available picornavirus-like polymerase sequences existed only as un-annotated transcripts within the transcriptomes of their hosts (Obbard 2018). Together, these two sources of (meta-)genomic data have ‘filled in’ the tree of viruses at many levels. They have expanded the host range of known viruses (e.g. Galbraith et al. 2018), identified vast numbers of likely new species and genera—consequently provoking considerable debate on how we should go about virus taxonomy (Simmonds et al. 2017, King et al. 2018, Simmonds and Aiewsakun 2018)—and identified new lineages that may warrant recognition at family level, including Chuviruses, Yueviruses, Qinviruses, Zhaoviruses, Yanviruses and Weiviruses (Li et al. 2015, Shi et al. 2016). More importantly, these discoveries have also started to impact upon our understanding of virus evolution (Wolf et al. 2018), emphasising the importance of ‘modular’ exchange (Koonin et al. 2015, Dolja and Koonin 2018) and suggesting surprisingly long-term fidelity to host lineages, at least at higher taxonomic levels (Geoghegan et al. 2017, Shi et al. 2018).

Despite the successes of metagenomic virus discovery, there are clear limitations to the approach. First, ‘virus-like sequences’ from a putative host need not equate to an active viral infection of that species. They may represent integrations into the host genome, infections of cellular parasites or other microbiota, infections of gut contents, or simply contaminating nucleic acid (reviewed in Obbard 2018). Second, most metagenomic methods rely on similarity searches to identify virus sequences through inferred homology. This limits the new discoveries to the relatives of known viruses. In the future, as similarity search algorithms become more sensitive (e.g. Kuchibhatla et al. 2014, Yutin et al. 2018), this approach may be able to uncover all viruses—at least those that have common ancestry with the references. However, this approach will probably still struggle to identify less conserved parts of the genome, especially for segmented viruses and incomplete assemblies. As a consequence, there may be many viruses and virus fragments that cannot be seen through the lens of homology-based metagenomics, the so-called ‘dark’ viruses (Rinke et al. 2013, Krishnamurthy and Wang 2017, Knox et al. 2018).

The ultimate solution to the shortcomings of metagenomic discovery is to isolate and experimentally characterise viruses. However, the large number of uncharacterised virus-like sequences means that this is unlikely to be an option in the foreseeable future. Instead, we can use other aspects of metagenomic data to corroborate evidence of a viral infection (reviewed in Obbard 2018). For example, metagenomic reads are more consistent with an active infection if RNA is very abundant (several percent of the total), if strand biases reflect active replication (such as the presence of the coding strand for negative sense RNA viruses or DNA viruses), or if RNA virus sequences are absent from DNA. The presence and absence of contigs across datasets can also provide useful clues as to the origin of a sequence. Specifically, sequences that are present in all individuals, or in all populations, are more likely to represent genome integrations, sequences that always co-occur with recognisable viral fragments may be segments that are not detectable by homology, and sequences that co-occur with non-host sequences are candidates to be viruses of the microbiota.

One of the most powerful ways to identify viruses is to capitalise on the host’s own ability to recognise pathogens, for example by sequencing the copious virus-derived small RNAs generated by the antiviral RNAi responses of plants, fungi, nematodes and arthropods (Aguiar et al. 2015, Webster et al. 2015). This not only demonstrates host recognition of the sequences as viral in origin, but also (if both strands of ssRNA viruses are present) demonstrates viral replication, and can even identify the true host of the virus based on the length distribution and base composition of the small RNAs (compare Webster et al. 2016, with Coyle et al. 2018).

Using ribosome-depleted RNA and small-RNA metagenomic sequencing, Webster *et al* (2015) previously proposed approximately 60 ‘dark’ virus sequences associated with *Drosophila.* These comprised contigs of at least one 1kbp that were present as RNA but not DNA, contained a long open reading frame, lacked identifiable homology with known viruses or cellular organisms, and were substantial sources of the 21nt small RNAs that characterise *Drosophila* antiviral RNAi. They included ‘Galbut virus’ (KP714100, KP714099), which has since been shown to constitute two divergent segments of an insect-infecting Partitivirus (KP757930; Shi et al. 2018) and is the most common virus associated with *Drosophila melanogaster* in the wild (Webster et al. 2015); ‘Chaq virus’ (KP714088), which may be a satellite or an optional segment of Galbut virus (Shi et al. 2018); and 56 unnamed ‘dark’ virus fragments (KP757937-KP757993). Subsequent discoveries have since allowed 26 of these previously dark sequences to be identified as segments or fragments of viruses that display detectable homology in other regions, including several pieces of Flavi-like and Ifla-like viruses (Shi et al. 2016, Shi et al. 2016) and the missing segments of a Phasmavirus (Matthew J. Ballinger, pers. com.) and Torrey Pines reovirus (Shi et al. 2018).

Here we combine data from Webster *et al* (2015) with a search of transcriptome assemblies and selected meta-transcriptomic datasets to identify six of the remaining ‘dark’ *Drosophila* virus sequences as segments of the founding members of a new lineage of segmented positive-sense single-stranded (+ss)RNA viruses. The protein encoded by segment 5 of these viruses shares a small number of conserved residues with the RNA dependent RNA polymerases of Picornaviruses, Flaviviruses, Permutotetraviruses, Reoviruses, Totiviruses and Picrobirna-viruses, but is not substantially more similar or robustly supported as sister to any of these lineages—suggesting that the new lineage may warrant recognition as a new family. We find at least one homologous segment in publicly-available transcriptomic data from each of 40 different animal species, including multiple arthropods and a small number of vertebrates, suggesting these viruses are associated with a diverse range of animal taxa.

## 2 Methods

### 2.1 Association of ‘dark’ virus segments from Drosophila

Webster *et al* (2015) previously performed metagenomic virus discovery by RNA sequencing from a large pool of wild-collected adult *Drosophila* (Drosophilidae; Diptera). In brief, *ca.* 5000 flies were collected in 2010 from Kenya (denoted pools E and K), the USA (pool I), and the UK (pools S and T). Ribosome depleted and double-stranded nuclease normalised libraries were sequenced using the Illumina platform, and assembled using Trinity (Grabherr et al. 2011). Small RNAs were sequenced from the same RNA pools, and the characteristic Dicer-mediated viral small-RNA signature used to identify around 60 putative ‘dark’ virus sequences that lacked detectable sequence homology (Supporting Figures S1 and S2; sequences accessions KP757937-KP757993). Raw data are available under NCBI project accession PRJNA277921. For details, see Webster *et al* (2015).

Here we took four approaches to identify sequences related to these ‘dark’ viruses of *Drosophila*, and to associate ‘dark’ fragments into viral genomes based on the co-occurrence of homologous sequences in other taxa. First, we obtained the collated transcriptome shotgun assemblies available from the European Nucleotide Archive (ftp://ftp.ebi.ac.uk/pub/databases/ena/tsa/public/; Most recently accessed 9^th^ August 2019) and inferred their protein sequences for similarity searching by translating all long open reading frames present in each contig. We used these to build a database for Diamond (Buchfink et al. 2014), and used Diamond ‘blastp’ to search the database with the translated ‘dark’ virus sequences identified from *Drosophila*. Second, we downloaded the pre-built tsa_nt BLAST database provided by NCBI (ftp://ftp.ncbi.nlm.nih.gov/blast/db/), and used tblastn (Camacho et al. 2009) to search this database for co-occurring homologous fragments with the same sequences. Third, we used diamond ‘blastx’ (Buchfink et al. 2014) to search large-scale metagenomic assemblies derived from various invertebrates (Shi et al. 2016) and vertebrates (Shi et al. 2018). For sources of raw data see Supporting File S1. Fourth, to identify missing fragments associated with *Drosophila*, we also re-queried translations of the raw unannotated meta-transcriptomic assemblies of Webster *et al* (2015) (https://doi.org/10.1371/journal.pbio.1002210.s002) using blastp (Camacho et al. 2009). Fragments with homologous sequences that consistently co-occurred across multiple transcriptomic datasets were taken forward as candidate segments of new viruses.

### 2.2 Identification of related viral segments from Lysiphlebus fabarum

Transcriptomic data were collected from adults and larvae of the parasitoid wasp *Lysiphlebus fabarum* (Braconidae; Hymenoptera) as part of an experimental evolution study (Dennis et al. 2017, Dennis et al. In Revision). Briefly, parasitoids were reared in different sublines of the aphid *Aphis fabae*, each either possessing different strains of the defensive symbiotic bacterium *Hamiltonella defensa*, or no *H. defensa*. Aphid hosts were reared on broad bean plants (*Vica faba*) and parasitoids were collected after 11 (adults) or 14 (larvae) generations of experimental selection. Poly-A enriched cDNA libraries were constructed using the Illumina TruSeq RNA kit (adults) or the Illumina TruSeq Stranded mRNA kit (larvae). Libraries were sequenced in single-end, 100bp cycles on an Illumina HiSeq2500 (sequence data available under NCBI PRJNA290156). Trimmed and quality filtered reads were assembled *de novo* using Trinity (v2.4.0, for details see Dennis et al. In Revision), read-counts were quantified by mapping to the reference using Bowtie2 (Langmead and Salzberg 2012), and uniquely-mapping read counts were extracted with eXpress (Roberts and Pachter 2012). To assign taxonomic origin, the assembled *L. fabarum* transcripts were used to query the NCBI *nr* protein blast database (blastx, e-values < 10^−10^). The subsequent differential expression analysis identified several highly expressed fragments that were not present in the *L. fabarum* draft genome nor in transcripts from the host aphid (*A. fabae*), and were not identified in the whole-transcriptome annotation using *blastn*. Subsequent protein-level searches (blastp, E-values < 10^−10^) revealed sequence similarity in four of the fragments to putative ‘dark’ virus sequences from *Drosophila* (Dennis et al. In Revision). Here we used read counts to confirm the co-occurrence of homologous fragments across *L. fabarum* individuals, and to identify a fifth viral segment, which had not previously detected on the basis of the original small RNA profile in *Drosophila*, on the basis of its co-occurrence across samples. To generate a complete viral genome, we selected a high-abundance larval dataset (ABD-118-118, SRA sample SAMN10024157, project PRJNA290156), for re-assembly with Trinity (Grabherr et al. 2011). For this assembly we subsampled the reads by 10 thousand-fold, as we have found that at very high levels of coverage, read-depth normalisation allows low-frequency polymorphisms to disrupt assemblies.

### 2.3 Determination of the genomic strand from a related virus of Lepidoptera

Strand-specific RNA libraries can be used to identify strand-biases in viral RNA, providing a clue as to the likely genomic strand of the virus and evidence for replication. Positive sense single-stranded (+ssRNA) viruses tend to be very strongly biased to positive sense reads, replicating double-stranded (dsRNA) viruses are weakly biased toward positive-sense reads, and replicating negative sense (-ssRNA) viruses are weakly biased toward negative sense reads. This is because mRNA-like expression products of replicating viruses have an abundance approaching that of the genomic strand. Unfortunately, much RNA sequencing is strand-agnostic (including that from the *Drosophila* datasets of Webster *et al* (2015)) and the vast majority of Eukaryotic transcriptomic datasets are sequenced from poly-A enriched RNA (such as that from *Lysiphlebus fabarum*), which artificially enriches for polyadenylated RNAs such as mRNA-like expression products. We therefore sought relatives in a strand-specific meta-transcriptomic dataset that had been prepared without poly-A enrichment.

For this purpose, we used a metagenomic dataset prepared as part of an ongoing study of British Lepidoptera (Longdon & Obbard, unpublished). Briefly, between one and twelve adults (total of 45) of each of 16 different species were collected from Penryn (Cornwall, UK) and Buckfastleigh (Devon,UK) in July and September 2017 respectively. Total RNA was extracted from each individual using Trizol-Chloroform extractions according to the manufacturer’s instructions, and a strand-specific library prepared from the combined pool using an Illumina TruSeq stranded total RNA kit treating samples with Gold rRNA removal mix. This was sequenced by the Exeter University Sequencing service using the Illumina platform. The reads were assembled *de novo* using Trinity (Grabherr et al. 2011), and the resulting assemblies searched as protein using Diamond ‘blastp’ (Buchfink et al. 2014).

We then used an RT-PCR screen to confirm the identity of the host, and to confirm that the five putative segments co-occurred in the same individual. RNA was reverse-transcribed using GoScript reverse transcriptase (Promega) with random hexamer primers, then diluted 1:10 with nuclease free water. PCRs to amplify short regions from the five viral segments (S1-S5) were carried out with the following primers: S1F ATGCATCTCGTTCCTGACCA and S1R GCCCCTTCAGACAGCTCTAA; S2F CACCACCAAGAACGGACAAG and S2R TGCCACCAC-TCTAACCACAT; S3F AGCAATTCAACGAC-CACACC and S3R GATAGGGGACAGGG-CAGATC; S4F ATGAACGAGAGGTGCCTTCA and S4R CTCCATCACCTTGACATGCG; S5F TGCACTGTTCAGCTACCTCA and S5R CCGTGTCGTTCGATGAAGTC, using a touch-down PCR cycle (95°C 30 sec, 62°C (−1°C per cycle) 30 sec, 72°C 1 min; for 10x cycles followed by; 95°C 30 sec, 52°C 30 sec, 72°C 1 min; for a further 30x cycles). As a positive control for RT we used host Cytochrome Oxidase I amplified with LCO/HCO primers (Folmer et al. 1994) (94°C 30 sec, 46°C 1 min, 72°C 1 min; for 5x cycles followed by; 94°C 30 sec, 50°C 1min, 72°C 1 min; for a further 35x cycles). All PCR reactions were carried out in duplicate using Taq DNA Polymerase and ThermoPol Buffer (New England Biolabs). We used (RT negative) PCR to confirm that none of these segments were present as DNA. To confirm the identity of the resulting PCR products, positive samples were Sanger sequenced from the reverse primer using BigDye (Applied Biosystems) after treatment with exonuclease I and shrimp alkaline phosphatase.

### 2.4 Inference of protein domain homology

Searches using blastp had previously been unable to detect homology between the putative ‘dark’ virus sequences of *Drosophila* and known proteins (Webster et al. 2015). However, more sophisticated Hidden Markov Model approaches to similarity searching that use position-specific scoring matrix (PSSM) profiles are known to be more sensitive (Kuchibhatla et al. 2014). We therefore aligned the putative viral proteins from *Drosophila* with their homologs from other transcriptomic datasets using MUSCLE (Edgar 2004), and used these alignments to query PDB, Pfam-A (v.32), NCBI Conserved Domain (v.3.16) and TIGRFAMs (v.15.0) databases using HHpred (Zimmermann et al. 2018).

### 2.5 Phylogenetic analysis

To infer relationships among the new viruses, we aligned protein sequences using Mcoffee from the T-coffee package (Wallace et al. 2006), and inferred relationships by maximum likelihood using IQtree (Nguyen et al. 2014). For each of the segments available from *Drosophila, L. fabarum, Lepidoptera*, and the other species, between 13 (Segment 3) and 41 (Segment 5) protein sequences were aligned, depending on level of sequence conservation. Regions of low conservation at either end of the alignments were selected by eye and removed. However, no internal regions were trimmed, as trimming leads to bias toward the guide tree and gives false confidence (Tan et al. 2015). The end-trimmed alignments were then used to infer phylogenetic relationships for each of the segments using the LG protein substitution matrix (Le and Gascuel 2008) with inferred residue frequencies and a 4-category discretised gamma distribution of rates.

To illustrate the relative distance (and likely unresolvable relationships) between the new viruses and previously described virus families, we selected for phylogenetic analysis the RNA dependent RNA polymerase (RdRp) sequences from representatives of major lineages of +ssRNA viruses. We aligned a core RdRp sequence of 215-513 residues for a total of 255 viruses, using two different methods; T-coffee ‘Expresso’ (Armougom et al. 2006), which uses structural data to inform the alignment, and T-coffee ‘accurate’, which combines structural data and protein profiles. Each of these alignments was used to infer the phylogenetic relationship of these clades by maximum likelihood, using IQtree as described above (Nguyen et al. 2014). As before, alignment ends were trimmed by eye, but not internally (Tan et al. 2015). To examine the consequences of conditioning on a specific alignment, we also inferred sequence relationships using BALi-Phy (Redelings 2014). BALi-Phy uses a Bayesian MCMC sampler to jointly infer the alignment, the tree, and the substitution and indel model parameters. Although computationally expensive, this captures some of the uncertainty inherent in inferring homology during alignment, and empirically BALi-Phy performs well with highly divergent sequences (Nute et al. 2018). We ran 22 simultaneous instances of BALi-Phy (totalling approximately 1.7 CPU years; Xeon E5-2620 v4 @2.10GHz), analysing the combined output after the effective sample size for most of the parameters (including the topological ESS) was in excess of 5000 and the potential scale reduction factor for these parameters less than 1.01. The exceptions were three parameters relating to the absolute evolutionary rate (tree scale) and the distribution of rates across sites. These occasionally flipped between two solutions with identical likelihoods, and had overall effective sample sizes of *ca.* 200. We do not believe this is likely to compromise our conclusions regarding the uncertainty in tree topology.

## 3 Results

### 3.1 Four segments of a ‘dark’ virus associated with Drosophila and other arthropods

We hypothesised that although the putative ‘dark’ virus fragments proposed by Webster *et al* (2015) on the basis of small-RNA profiles (Supporting Figures S1 and S2) lacked detectable homology with known viruses, their relatives may be present—but unrecognised—in transcriptome assemblies from other species. If so, we reasoned that the co-occurrence of homologous sequences across different datasets could allow fragments from *Drosophila* to be associated into complete virus genomes. Using similarity searches we initially identified six fragments from Webster *et al* (2015) that each consistently identified homologs in several distantly related transcriptomic datasets; those of the centipede *Lithobius forficatus* (transcriptome GBKE; NCBI project PRJNA198080 (Rehm et al. 2014)), the locust *Locusta migratoria manilensis* (GDIO; PRJNA283919 (Zhang et al. 2015)), the leafhopper *Clastoptera arizonana* (GEDC; PRJNA303152 (Tassone et al. 2017)), the hematophagous bug *Triatoma infestans* (GFMC; PRJNA304741 (Traverso et al. 2017)), and two parasitoid wasps, *Ceraphron* spp. (GBVD; PRJNA252127 (Peters et al. 2017)) and *Psyttalia concolor* (GCDX; PRJNA262710). Motivated by this discovery of four homologous sequence groups across these taxa, we performed a new search of the Webster *et al* (2015) data that identified two additional fragments. The eight *Drosophila*-associated sequences formed two groups (four sequences from drosophilid pool E and four from drosophilid pool I) encoding proteins that ranged between 40% and 60% amino acid identity (See supporting File S1 for accession numbers).

Subsequent searches later identified homologs in 14 other arthropod transcriptomes, including six from Hymenoptera, five from Hemiptera, two from Coleoptera, and one each from Lepidoptera and Odonata (Supporting File S1). We also identified some segments in a plant transcriptome (*Jasminum sambac*; PRJNA551353; SAMN12158026). However, as this dataset contained a large number of reads from the Jasmine whitefly *Dialeurodes kirkaldyi*, we think it unlikely that the plant is the true host.

Although none of the protein sequences from these fragments displayed significant blastp similarity to characterised proteins, the presence of the four clear homologs in eight unrelated arthropod transcriptomes strongly supported an association between them. In addition, the similar length and similar coding structure of the fragments across species suggested that they comprise the genomic sequences of a segmented virus (all between 1.5 and 1.7 kbp, containing a single open reading frame; Figure 1). Finally, as expected for viruses of *Drosophila*, all segments were sources of 21nt small RNAs from along the length of both strands of the virus, demonstrating that the virus is recognised as a double-stranded target by Dicer-2 (Supporting Figures S1 and S2). We therefore speculatively named these putative viruses from drosophilid pools E and I as ‘Kwi virus’ and ‘Nai virus’ respectively, and submitted them to GenBank (KY634875-KY634878; KY634871-KY634874; mentioned in Obbard 2018). Provisional names were chosen following the precedent set by *Drosophila* ‘Nora’ virus (*new* in Armenian (Habayeb et al. 2006)) and ‘Galbut’ virus (*maybe* in Lithuanian (Webster et al. 2015)), with *Kwí* and *Nai* being indicators of uncertainty (*maybe, perhaps*) in JRR Tolkien’s invented language Quenya (Wickmark 2019).

**Figure 1:**
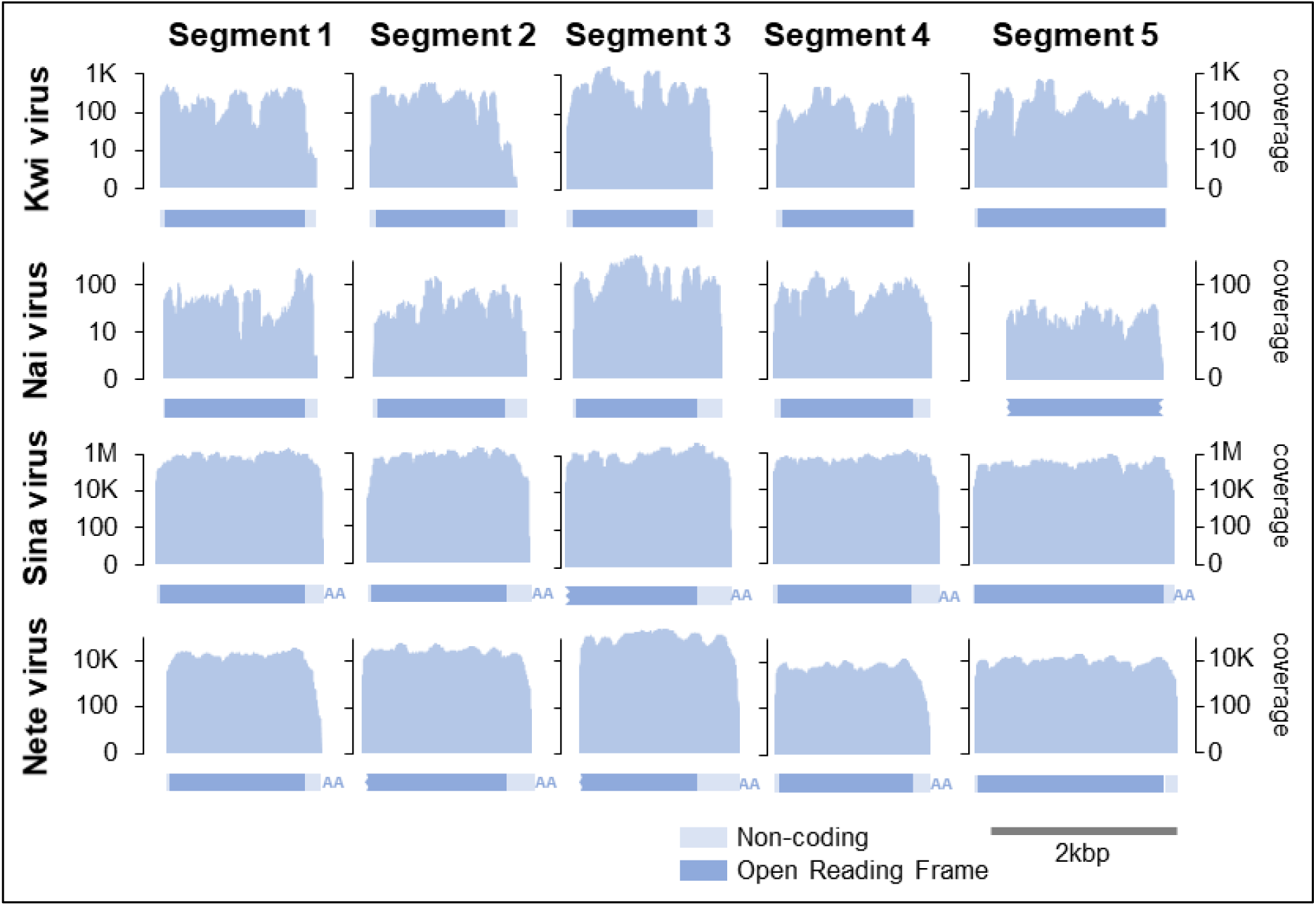
Virus segments and sequencing coverage. Panels show the structure and fold-coverage for each of the five segments (columns), for each of the four founding Quenyaviruses (rows). Graphs represent fold-coverage on a log_10_ scale, with the structure of the segment annotated below to scale (dark: coding, pale: non-coding). Assembled contigs that terminated with a poly-A tract are denoted ‘AA’), and potentially incomplete open reading frames indicated with a jagged edge.

### 3.2 A related hymenopteran virus identifies a fifth segment

In an unrelated expression study of the parasitoid wasp *Lysiphlebus fabarum*, Dennis *et al* (In Revision) identified four sequences showing clear 1:1 homology with the segments of Kwi virus and Nai virus. These were again *ca.* 1.5kb in length, and each encoded a single open reading frame (Figure 1). Each segment had a poly-A tract at the 3’ end, suggesting either that the virus has poly-adenylated genome segments, or that these represent poly-adenylated mRNA-like expression products. Strongly consistent with a viral origin, the sequences were present in some individuals but not others (Supporting Figure S3), always co-occurred with correlated read numbers (correlation coefficient >0.87; Supporting Figure S3C), and could be extremely abundant—accounting for up to 40% of non-ribosomal reads and equating to 1 million-fold coverage of the virus in some wasps (Figure 1).

Based on the high abundance and the clear pattern of co-occurrence, we searched for other wasp-associated contigs displaying the same properties, reasoning that these were likely to be additional segments of the same virus. This search identified a candidate 5^th^ segment of *ca.* 2kbp, again containing a single open reading frame (Figure 1). We then sought homologs of this 5^th^ segment in the data of Webster *et al* (2015) and in the public transcriptomic datasets outlined above. As expected, we were able to find a homolog in almost every case, confirming co-occurrence of the five putative viral segments across datasets (Figure 1; supporting File S1; Nai virus NCBI accession MH937729, Kwi virus MH937728). The protein encoded by the newly-identified segment 5 was substantially more conserved than the other proteins, with 64% amino-acid identity between Kwi virus and Nai virus. We believe that it had most likely been missed from the putative ‘dark’ viruses of Webster *et al* (2015) because of the relatively small number of reads present in that dataset (10-100 fold coverage; Figure 1). Based on these segments, we used a re-assembly of a single larval *Lysiphlebus fabarum* dataset (sample ABD-118; Supporting Figure 3) to provide an improved assembly, which we provisionally named ‘Sina Virus’, reflecting our increased confidence that the sequences are viral in origin (*Sína* is Quenya for *known, certain, ascertained*) and submitted the sequences to Genbank under accession numbers MN264686-MN264690.

### 3.3 A related Lepidopteran virus suggests +ssRNA as the genomic material

To determine whether these virus genomes are likely to be double-stranded RNA (dsRNA), positive sense single stranded (+ssRNA) or negative sense single-stranded (-ssRNA), we identified a related virus in a strand-specific metatranscriptomic dataset that had been prepared without poly-A enrichment from several species of Lepidoptera (Longdon & Obbard, unpublished). All 5 segments were detected (Figure 1), and as was the case for Kwi, Nai, and Sina viruses, segments 1-4 were around 1.6kbp and segment 5 around 2kbp in length, each containing a single open reading frame (Figure 1). We have provisionally named these sequences as ‘Nete virus’ (*Netë* is Quenya for *another one, one more*) and submitted them to GenBank under accession numbers MN264681-MN264685.

Overall, this virus accounted for 3% of the reads in the metagenomic pool, giving around 10 thousand-fold coverage of the genome (Figure 1). An analysis of the strand bias in the metagenomic sequencing found that 99.8% of reads derived from the positive-sense (coding) strand, strongly suggesting that this virus has a +ssRNA genome (Supporting File S2). The five segments appeared complete at the 3’ end, possessing a poly-A tail and suggesting that the genomic +ssRNA is polyadenylated (Figure 1). Four of the five segments (excluding segment 2) possessed a conserved sequence of *ca.* 150nt at the 3’ end, and a similar pattern (but not sequence) was seen in the closely related segments from the *Ceraphron sp.* transcriptome. However, we were unable to identify any 5’ pattern or motif shared among the segments.

An RT-PCR survey of the individual moth RNA extractions used to create the metagenomic pool showed that all five segments co-occur in a single *Crocallis elinguaria* individual (Geometridae; Lepidoptera), collected at latitude 50.169, longitude −5.125 on 23/Jul/2017. RT-negative PCR showed that viral segments were not present in a DNA form.

### 3.4 Related viruses are present in metagenomic datasets from other animals

After identifying the complete (five segment) virus genomes in transcriptomic datasets from 12 different arthropods, and incomplete genomes (between one and four segments) in a further 15 arthropod datasets (Supporting File S1), we sought to capitalise on recent metagenomic datasets to identify related sequences in other animals (Shi et al. 2016, Shi et al. 2018). This search yielded complete (or near-complete) homologs of segment 5 (the most conserved protein) in 18 further datasets, including four from mixed pools of insects, two from spiders, three from crustaceans, seven from bony fish, and one each from a toad (Dongxihu virus associated with *Bufo gargarizans*) and a lizard (Bawangfen virus associated with *Calotes versicolor*). Five of these pools also contained homologs of segment 1 (the second most conserved protein), and one also contained segment 4 (the third most conserved protein). These sequences have been submitted to Genbank under accession identifiers MN371231-MN371254; See Supporting File S1 for details.

The finding that these virus sequences can be associated with both vertebrates and invertebrates may indicate that they are broadly distributed across the metazoa (note the only non-metazoan associated sequence came from a plant trascriptome contaminated with insects). However, metagenomic data alone cannot confirm this, as such datasets can include contamination from gut contents or parasites of the supposed host taxon. We therefore explored four sources of evidence that could be used to corroborate the targeted taxon as the true host. First, we examined the viral read abundance, as very high abundance is unlikely for viruses of contaminating organisms. Abundance ranged from over 37,124 Reads Per Kilobase per Million reads (RPKM; 40% of non-ribosomal RNA) for Sina virus in one *Lysiphlebus fabarum* sample, to 0.16 RPKM (six read-pairs) for Zhanggezhuang virus from a metagenomic pool of Branchiopoda, with a median of 16.9 RPKM (Supporting File S1). This strongly supports some of the arthropods (such as *Lysiphlebus*) as true hosts, but does not support or refute that the virus may infect vertebrates (e.g. RPKM as high as 834 for one Scorpaeniformes fish sample, but as low as 4.6 in *Drosophila* Nai virus, where infection could be independently confirmed by the presence of 21nt viral small RNAs). Second, for two high-coverage low species-complexity vertebrate metagenomic pools (the *Bufo gargarizans* lung sample and *Calotes versicolor* gut sample) we searched raw assemblies for Cytochome Oxidase I sequences of contaminating invertebrates. This found that <0.5% of the RNA from *Bufo gargarizans* (Dongxihu virus RPKM of 94.6) and <0.01% of RNA from *Calotes versicolor* (Bawangfen virus RPKM of 338.5) derived from contaminating invertebrates, strengthening the possibility that the vertebrate is the true host. Third, for Segment 5 (which was available for most taxa) we examined the deviation in dinucleotide composition from that expected on the basis of the base composition, as this is reported to be predictive of host lineages (Kapoor et al. 2010, but see Di Giallonardo et al. 2017). However, we were unable to detect any clear pattern among viruses, either by inspection of a PCA, or using a linear discriminant function analysis. This may support a homogenous pool of true hosts, such as arthropods but not vertebrates, but the short sequence length available (<2kbp) and small sample size (32 sequences) means that such an analysis probably lacks power.

Finally, we also analysed the phylogenetic relationships for all of the segments, as (except for vectored viruses) transitions between vertebrate and invertebrate hosts are generally rare (Longdon et al. 2015, Geoghegan et al. 2017). This showed that, despite the apparent absence of contaminating invertebrates, sequences from the toad (Dongxihu virus) and the lizard (Bawangfen virus) both sit among arthropod samples (segments 1 and 5; Figure 2E), as do the several other sequences from fish. The analysis also identified a deeply divergent clade of four sequences from bony fish with no close relatives in invertebrates that, if not contamination, could in principle represent a clade of vertebrate-infecting viruses (Figure 2E). Accession numbers, alignments and tree files are provided in Supporting file S3.

**Figure 2:**
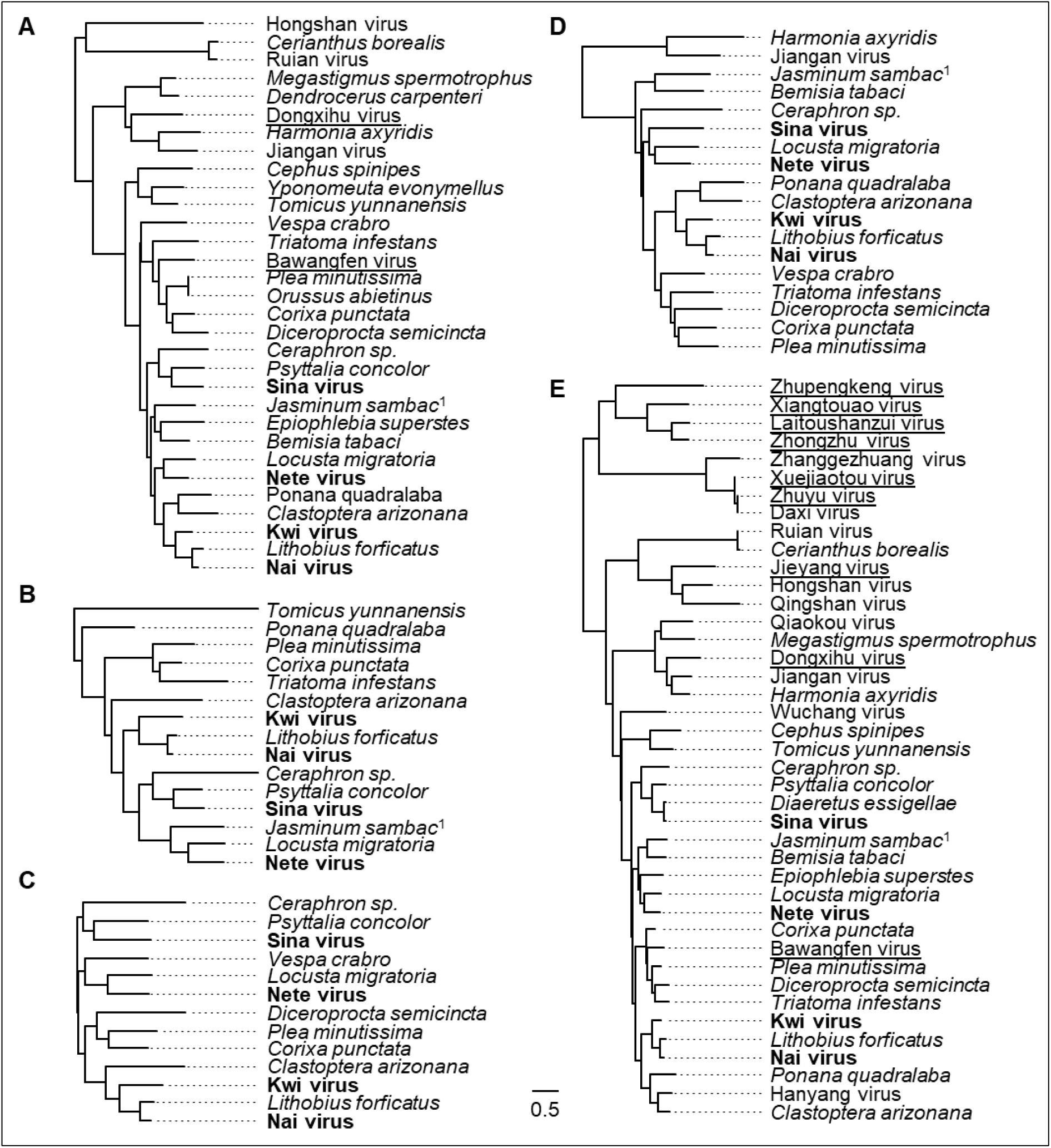
Phylogenetic trees for each of the viral segments. Panels A-E show maximum-likelihood phylogenetic trees for segments 1-5, inferred from amino-acid sequences. Panel E shows the tree for the most conserved segment, which encodes a putative RNA dependent RNA polymerase. Trees are mid-point rooted, and the scale bar represents 0.5 substitutions per site. The four viruses marked in bold are the founding members of the clade, those underlined come from nominally vertebrate metagenomic datasets, and species names in italic denote sequences from public transcriptomes. One, *Jasminum sambac* (marked superscript 1), came from a plant transcriptome contaminated with the whitefly *Dialeurodes kirkaldyi*. Note that some aspects of tree topology appear to be consistent among segments, suggesting that reassortment may be limited. Sequence alignments and tree files are provided in Supporting File S3

**Figure 3:**
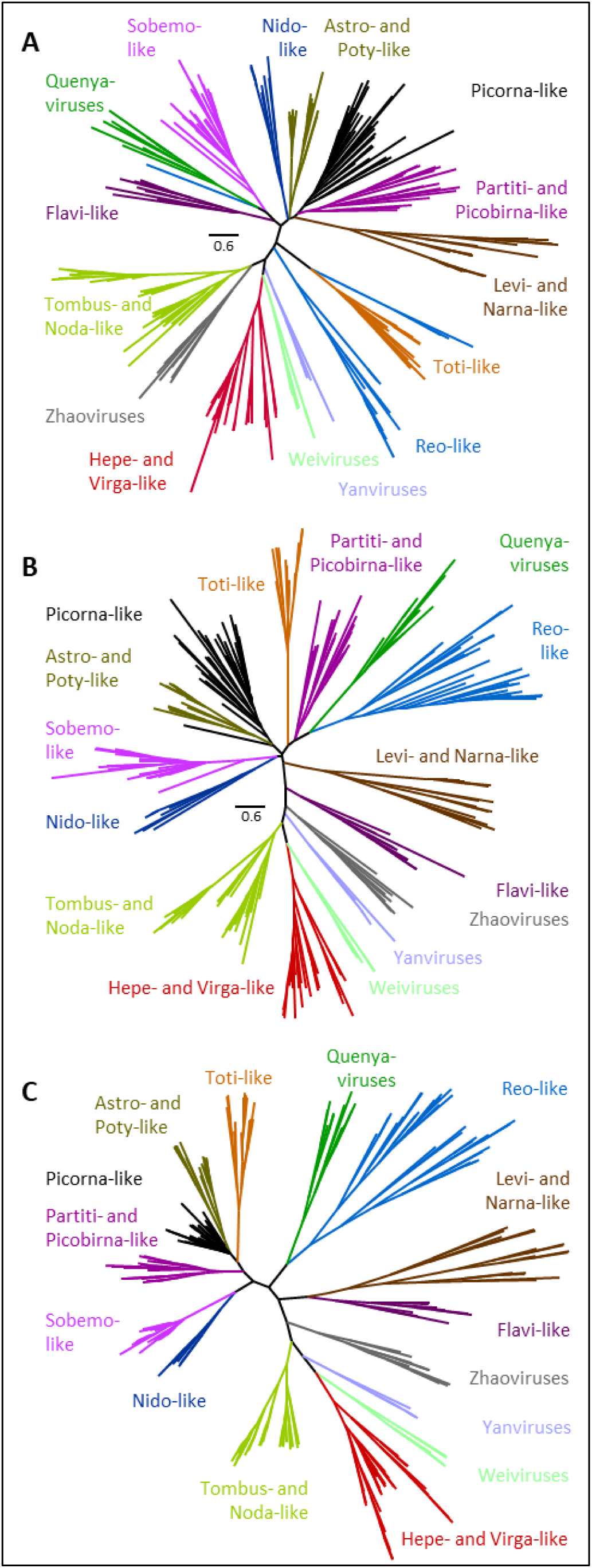
Relationship of the Quenyaviruses to other RNA viruses. Unrooted phylogenetic trees showing the possible relationships between the RdRp (segment 5) of Quenyaviruses and RdRps of representatives from other groups of RNA viruses. Trees were inferred by maximum-likelihood using IQtree from alignments using TCoffee modes ‘expresso’ (A) and ‘accurate’ (B), or using a Bayesian approach (C) that coinfers the tree and alignment. None of the deep relationships had any support in the Bayesian analysis, although all of the major clades were recovered and many of the relationships between them are the same as those in (B). Sequence alignments are provided in Supporting File S3.

### 3.5 Segment 5 has similarity to viral RNA dependent RNA polymerases

Having identified 1:1 homologs in multiple datasets, we were able to use the aligned protein sequences to perform a more sensitive homology search for conserved protein motifs using HHpred (Zimmermann et al. 2018). This still identified no significant similarity in the proteins encoded by segments 2 to 4 (E-value >1), and only a weakly-supported *ca.* 110 amino acid region of the segment 1 alignment with similarity to methyl-transferase / mRNA capping enzymes (E-value = 0.0019; see Supporting File S4). However, in contrast to searches using blastp, the alignment of segment 5 displayed a more strongly supported *ca.* 300 amino acid region with similarity to the RNA dependent RNA polymerase of Norwalk virus (E-value = 2.2×10^−33^; see Supporting File S4). This sequence was approximately equally matched to around 25 different reference structure or motifs, including RdRps from both +ssRNA viruses such as Picornavirales, Flavi-like viruses, and Permutotetraviruses, and dsRNA viruses such as Reoviruses, Picobirnaviruses, and Totiviruses. Notably, this region of similarity included a very highly conserved GDD motif that is shared by many viral polymerases, supporting the idea that segment 5 encodes the viral polymerase.

### 3.6 ‘Quenyaviruses’ are highly divergent and may constitute a new family

The new virus lineage described here has a distinctive genome structure comprising four 1.6kbp +ssRNA segments each encoding a single protein of unknown function, and one 2kbp +ssRNA segment encoding an RdRp. The putative RdRp is substantially divergent from those of characterised +ssRNA and dsRNA virus families, to the extent that similarity cannot be detected using routine blastp. On this basis we propose the informal name ‘Quenyaviruses’, reflecting the naming of the four founding members, and suggest that they may warrant consideration as a new unplaced family.

To explore their relationships with other RNA viruses using an explicit phylogenetic analysis, we selected a region of 215-513 amino acid residues of the core RdRp region from 11 representative Quenyaviruses and 244 other +ssRNA viruses, representing most major lineages. We excluded birnaviruses and permutotetraviruses, which have a permuted RdRp that cannot be straightforwardly aligned (Wolf et al. 2018). Phylogenetic inference is necessarily challenging with such high levels of divergence (mean pairwise protein identity of only 7.6%) and the inferred relationships among such distantly-related lineages are unlikely to be reliable (Bhardwaj et al. 2012, Nute et al. 2018). In particular, although current phylogenetic methods perform surprisingly well on simulated data with identities as low 8-10%, this is only true when homology is known (i.e. the true alignment is available; Bhardwaj et al. 2012). When the alignment has to be inferred, performance is poor—even when the true substitution model is the one being modelled (Nute et al. 2018). We therefore compared between trees that conditioned on each of two different alignment methods (Mcoffee modes ‘expresso’ and ‘accurate’), and also co-inferred the tree and the alignment using BALi-Phy (Redelings 2014). Accession numbers, alignments and tree files are provided in Supporting File S3.

All methods found the Quenyavirus RdRps to form a monophyletic clade, supporting their treatment as a natural group. Two of the methods placed the Quenyaviruses closer to (some of) the Reo-like viruses than to others (Figure 3B and C). However, there was little consistency in the placement of the other clades relative to each other. Moreover, the Bayesian joint alignment/tree analysis gave almost no posterior support to any of the major clades (Figure 3C; Supporting File S3). It is notable that many deep divisions seen in our three different approaches differ to those in the tree inferred by Wolf et al (2018), using maximum likelihood conditioned from an alignment in which sites with >50% gaps had been deleted. We believe that this suggests the relationships among these lineages cannot currently be robustly inferred. Nevertheless, the uncertainty in the placement of the Quenyaviruses emphasises their deep divergence from other taxonomically-recognised virus clades.

## 4. Discussion

Here we report the discovery of the Quenya-viruses, a new clade of segmented +ssRNA viruses identifiable from multiple (meta-)transcriptomic datasets, primarily of arthropods. Four of these segments had initially been identified as ‘dark’ viruses of *Drosophila*, purely on the basis of the characteristic small-RNA signature created by the host antiviral RNAi pathway (Webster et al. 2015). Now, by identifying a fifth segment encoding a divergent RdRp, we show that they form a monophyletic clade that is only distantly related to other +ssRNA viruses, and cannot be confidently placed within a wider phylogeny.

As with other metagenomic studies of virus diversity, this work raises two important questions. First, how well have we truly sampled the virosphere? Metagenomic studies often contain sequences lacking detectable homology, and it has been suggested that these include many ‘dark’ viruses (Krishnamurthy and Wang 2017). This may imply that many deeply-divergent viruses, or viruses lacking common ancestry with known families, remain to be discovered. Alternatively, many of the ‘dark’ sequences may be the less-conserved fragments of otherwise easily-recognised virus lineages (e.g. François et al. 2018). Thus far, of the predicted ‘dark’ *Drosophila* virus sequences of Webster *et al* (2015), 46% remain dark, 44% are now recognisable as members of known virus lineages, and 10% represent a genuinely new divergent lineage (the Quenyaviruses)—albeit one for which a sensitive search can identify some evidence of homology. Second, how many viruses are hiding in plain sight? Perhaps 10% of polymerase sequences from Picornavirales are currently unannotated as such within transcriptomic datasets (Obbard 2018), and surveys of publicly available data often identify multiple new viruses (e.g. François et al. 2016, Gilbert et al. 2019). Some of the sequences we analyse here have been in the public domain for more than 7 years, but without routine screening and annotation (or submission of such sequences to databases) they not only remain unavailable for analysis, but also potentially ‘contaminate’ other analyses with misattributed taxonomic information. Finally, our work also emphasises the ease with which new viruses can be identified, relative to the investment required to understand their biology. The Quenyaviruses seem broadly distributed, if not common, but we have no knowledge of their host range, transmission routes, tissue tropisms, or pathology.

## Supporting information

Accesstion numbers and contig properties

Strand bias in sequencing

Tree files and alignments, with accession numbers

HHpred analysis raw output

## Acknowledgements

We thank Christoph Vorburger for facilitating the re-use of data from *Lysiphlebus*, David Karlin for advice on the identification of protein domains, Benjamin Redelings for advice on the use of Bali-Phy, and all of the many researchers who provided their data to publicly available databases.

---

**Supporting Figure S1:**
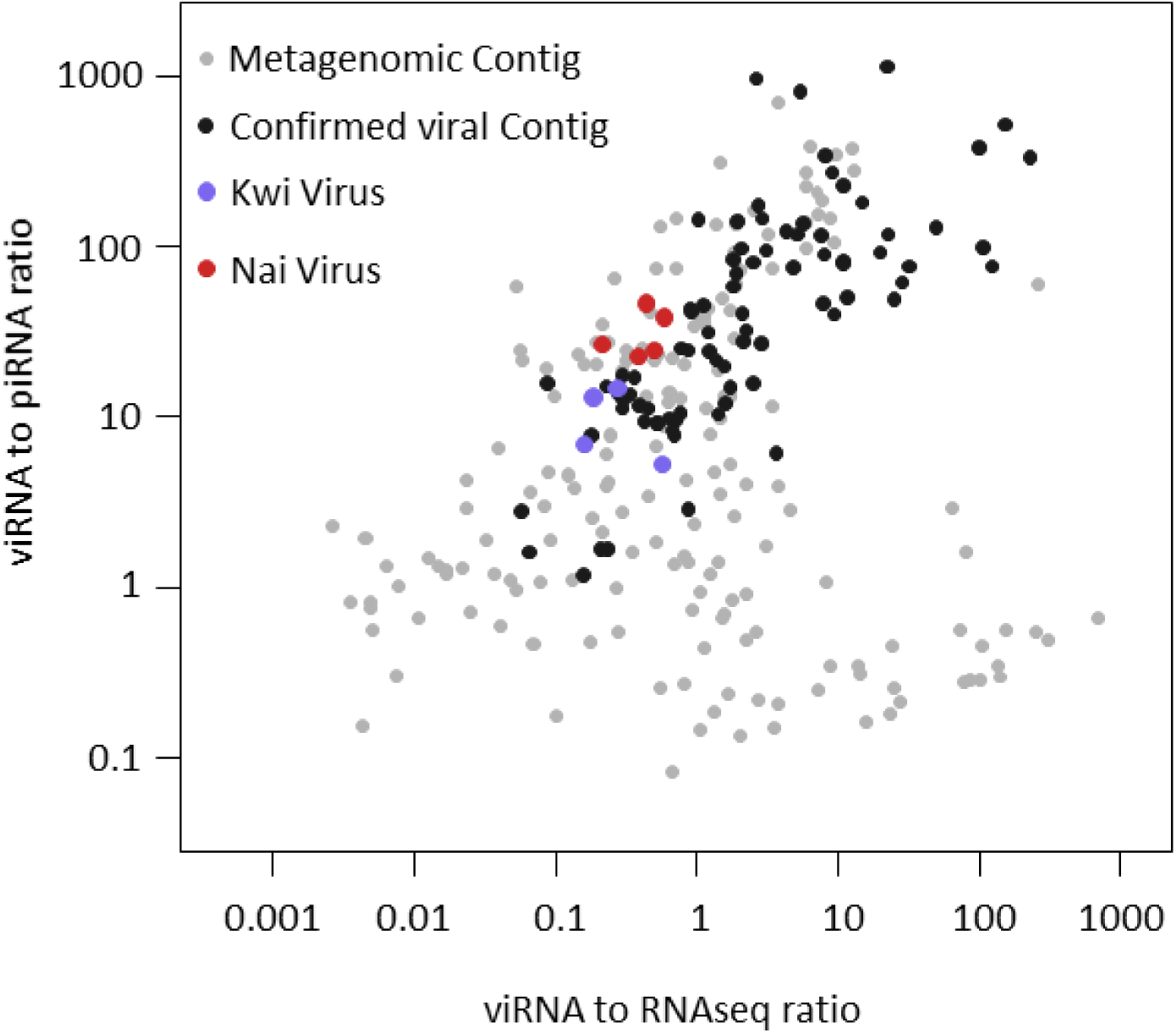
‘Dark’ virus identification by small-RNA sequencing. Points correspond to the contigs assembled by Webster et al (2015) that are sources of substantial numbers of small RNAs, and thus candidates to be viruses (high viRNA:piRNA length ratio) or transposable elements (low viRNA:piRNA ratio). Those marked in black have high blastp-detectable sequence similarity to known viruses, and those marked in colour correspond to segments of Kwi and Nai virus. Many pale grey points in the top-right corner of the plot are the other unconfirmed siRNA ‘candidate’ viruses reported by Webster et al (2015).

**Supporting Figure S2:**
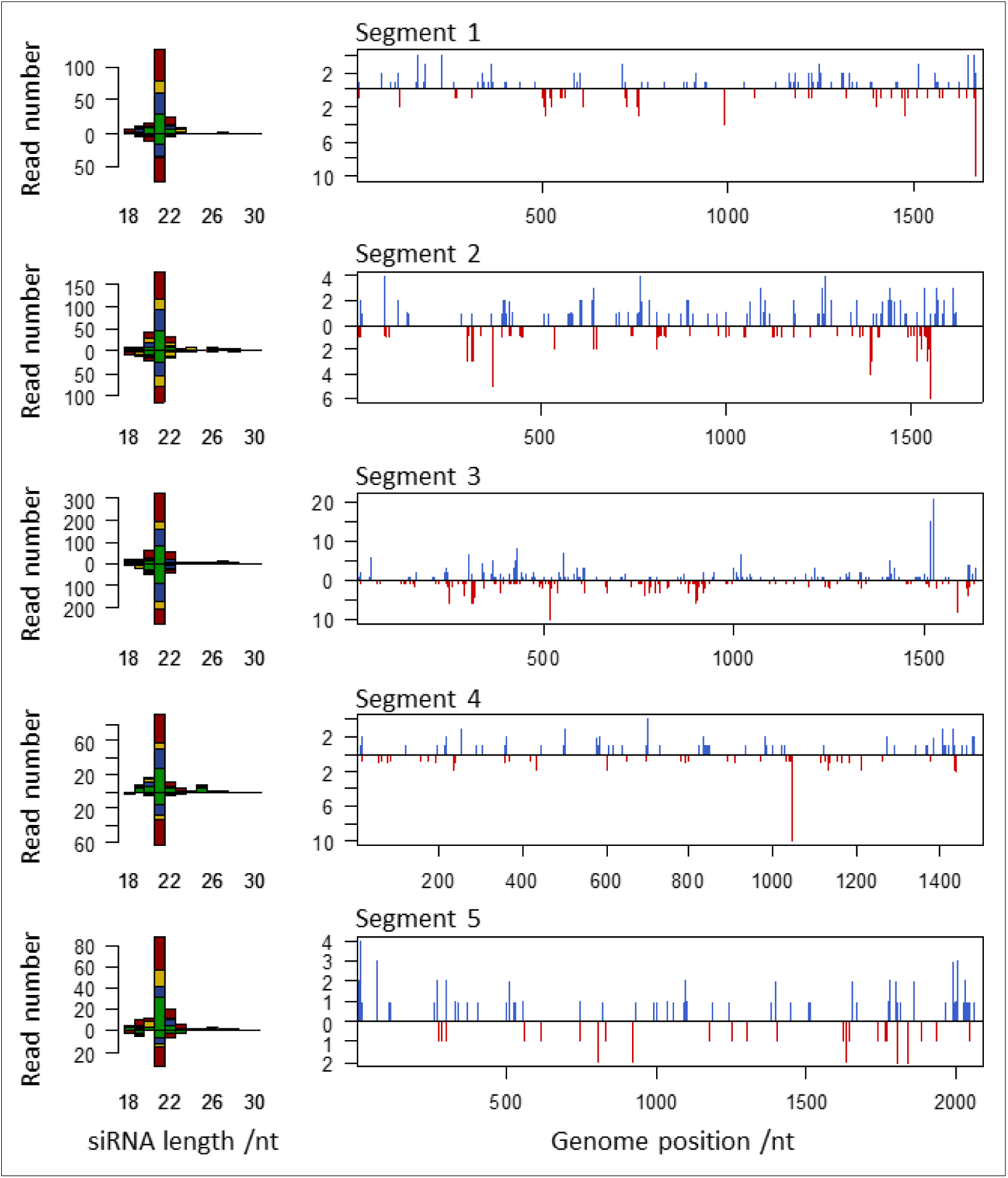
Kwi virus small RNA size distribution. The bar plots (left column) show the size distribution of reads mapping to each segment (rows 1-5) of Kwi virus. Bars are coloured according to the 5’ base (red U, yellow G, blue C and green A), numbers plotted above the x-axis show read counts mapping to the positive strand, and those below the x-axis mapping to the negative strand. Line plots (right column) show the genomic locations and numbers of the 21nt reads deriving from the positive (blue) and negative (red) strands of the virus. Note that siRNA numbers reflect the apparent abundance of each segment in other hosts (Supporting Figure S3).

**Supporting Figure S3:**
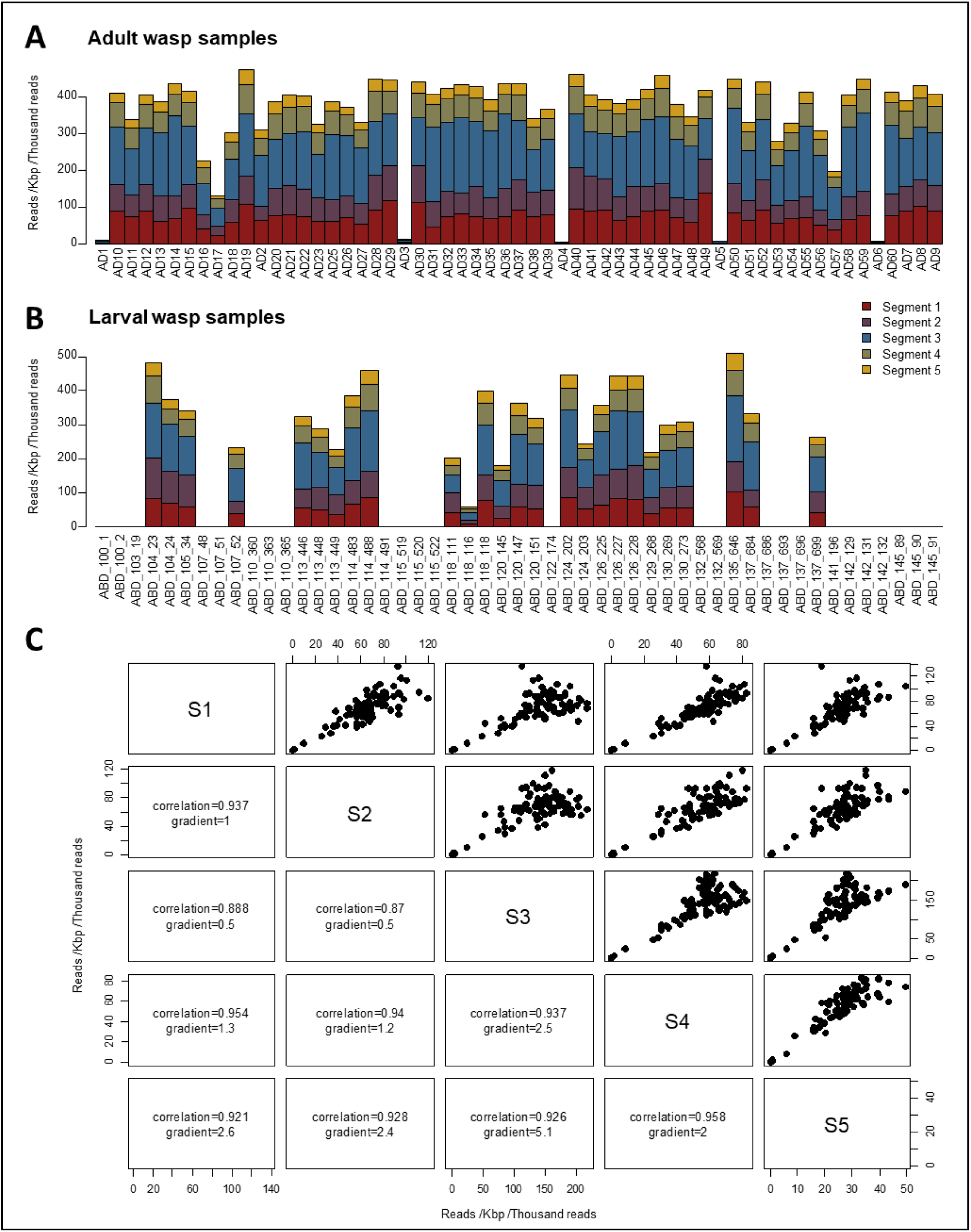
Co-occurrence of Sina virus segments across *L. fabarum* samples. Panels show the virus read abundance for each segment (colours) from each of the adult samples (A) and larval samples (B), and the correlation in read abundance between segments across all samples (C) on a scale of virus reads per kilobase per thousand total reads. Note that virus read numbers are highly correlated among segments (Panel C: correlation coefficient >0.87), and that reads from segment 3 are always most abundant while those from segment 5 are always least abundant (panel C). Note that Adult samples 1-3 were from the same experimental cage, as were 4-6.

**Supporting File S1: Virus details**

Excel table providing host species, NCBI project accessions, NCBI Samples, Read abundance and sequence accessions

**Supporting File S2: Strand bias in the sequencing reads from Lepidoptera**

Excel table giving the number of positive and negative sense forward-reads for each segment of Nete virus, with comparison ratios for high abundance viruses reported in Waldron et al (2018) and Medd et al (2018)

**Supporting File S3: Phylogenetic Analyses**

Compressed text files containing the alignments and tree files for Figures 2 and 3

**Supporting File S4: Raw HHpred output**

Compressed text files containing the raw output from the HHpred analysis

